# GenomeScope: Fast reference-free genome profiling from short reads

**DOI:** 10.1101/075978

**Authors:** Gregory W. Vurture, Fritz J. Sedlazeck, Maria Nattestad, Charles J. Underwood, Han Fang, James Gurtowski, Michael C. Schatz

## Abstract

**Summary:** GenomeScope is an open-source web tool to rapidly estimate the overall characteristics of a genome, including genome size, heterozygosity rate, and repeat content from unprocessed short reads. These features are essential for studying genome evolution, and help to choose parameters for downstream analysis. We demonstrate its accuracy on 324 simulated and 16 real datasets with a wide range in genome sizes, heterozygosity levels, and error rates.

**Availability and Implementation:** http://genomescope.org, https://github.com/schatzlab/genomescope.git

**Supplementary information:** Supplementary data are available at *Bioinformatics* online.

## 1 Introduction

High throughput sequencing enables the sequencing of novel genomes on a daily basis. However, even the most basic characteristics of these genomes, such as their size or heterozygosity rate, may be initially unknown, making it difficult to select appropriate analysis methods e.g. read mapper, *de novo* assembler, or SNP caller (Smolka, et al., 2015). Determining these characteristics in advance can reveal if an analysis is not capturing the full complexity of the genome, such as underreporting the number of variants or failure to assemble a significant fraction of the genome. Experimental methods are available for measuring some of these properties, although can require significant cost and labor.

A few computational approaches are now available for estimating the genome size from unassembled sequencing reads (Chikhi and Medvedev, 2014; Melsted and Halldorsson, 2014) and some genome assemblers internally compute related statistics to guide the algorithm (Bankevich, et al., 2012; Gnerre, et al., 2011). These methods follow earlier work to infer the length of BAC sequences from shotgun Sanger sequencing that analyze the frequency of sequences in the reads (Li and Waterman, 2003). However, only a few methods are available for measuring more complex characteristics such as the rate of heterozygosity and these methods can be difficult to use or interpret. Simpson (2014) proposed a computational method to estimate some of these properties from sequencing reads using *de novo* assembly techniques. However, this method is computationally intensive and can be difficult to interpret as results are reported relative to the assembly graph, such as the variant-induced branch rate rather than the more direct rate of heterozygosity. The GCE method (Liu, et al., 2013) also attempts to determine genome size and heterozygosity rate, but is not fully automated and requires users to manually specify several cutoffs. It also uses a Poisson coverage model that can lead to poor estimates with real sequencing data, and the heterozygosity model is limited to genomes without repetitive sequences.

## 2 Methods

Here we introduce *GenomeScope* to estimate the overall genome characteristics (total and haploid genome length, percentage of repetitive content, and heterozygosity rate) as well as overall read characteristics (read coverage, read duplication, and error rate) from raw short read sequencing data. The estimates do not require a reference genome and they can be automatically inferred via a statistical analysis of the *k-mer profile.* The *k-mer* profile (sometimes called *k-mer spectrum*) measures how often *k-mers*, substrings of length *k*, occur in the sequencing reads and can be computed quickly using tools such as Jellyfish (Marcais and Kingsford, 2011) or approximated using faster streaming approaches (Melsted and Halldorsson, 2014). The profiles reflect the complexity of the genome: homozygous genomes have a simple Poisson profile while heterozygous ones have a characteristic bimodal profile (Kajitani, et al., 2014). Repeats add additional peaks at even higher *k-mer* frequencies, while sequencing errors and read duplications distort the profiles with low frequency false *k-mers* and increased variances (Kelley, et al., 2010; Miller, et al.,2011).

Aware of these possible complexities, *GenomeScope* fits a mixture model of four evenly spaced negative binomial distributions to the *k-mer* profile to measure the relative abundances of heterozygous and homozygous, unique and two-copy sequences **(Supplemental Eq. 2)**. GenomeScope uses a mixture model of negative binomial model terms rather than Poisson terms since real sequencing data is often over-dispersed compared to a Poisson distribution (Miller, et al., 2011). The model fitting is computed using a non-linear least squares estimate as implemented by the *nls* function in R (Bates and Watts, 1988). To make the model fitting more robust, GenomeScope attempts several rounds of model fitting excluding different fractions of low frequency *k-mers* that are likely caused by sequencing errors, and adjusting for the ambiguity in determining the correct heterozygous and homozygous peak. The final set of parameters is selected as those parameters that minimize the residual sum of squares errors (RSSE) of the model relative to the observed *k-mer* profile. Afterwards, sequence errors and higher copy repeats are identified by *k-mers* falling outside the model range, and the total genome size is estimated by normalizing the observed *k-mer* frequencies to the average coverage value for homozygous sequences, excluding likely sequencing errors. See **Supplementary Note 1** for a detailed description of the model and fitting procedure.

GenomeScope is available open-source as a command line R application and also as an easy-to-use web application. Either version has minimum user requirements, consisting of (1) a text file of the k-mer profile computed by Jellyfish or other tools, (2) the value used for *k*, and (3) the length of the sequencing reads. Either the command line or online version of GenomeScope typically completes in less than 1 minute with modest RAM requirements, and outputs publication quality figures as well as text files with the inferred genome properties. If the modeling fails to converge, typically because of low coverage or low quality reads, the k-mer profile is plotted without the model parameters displayed so users can inspect the likely causes.

## 3 Results

We first applied GenomeScope to analyze 324 simulated data sets varying in heterozygosity (0.1%, 1%, 2%), average rate of read duplication (1, 2, 3), sequencing error rate (0.1%, 1%, 2%), coverage (100x, 50x, 25x, 15x) and organism (*E.coli, A. thaliana*, *D. melanogaster*) (**Supplementary Table 3, Supplementary Note 2**). A subset of the results for *A. thaliana* are displayed in **Figure 1A** (left), and show that the GenomeScope results are highly concordant with the true simulated rates over many conditions. The results were also highly concordant to a standard short-read variant analysis pipeline using BWA-MEM (Li, 2013) and SAMTools (Li, et al., 2009) or through whole genome alignment using DnaDiff (Phillippy, et al., 2008) of the original and mutated reference sequence.

**Figure 1.**
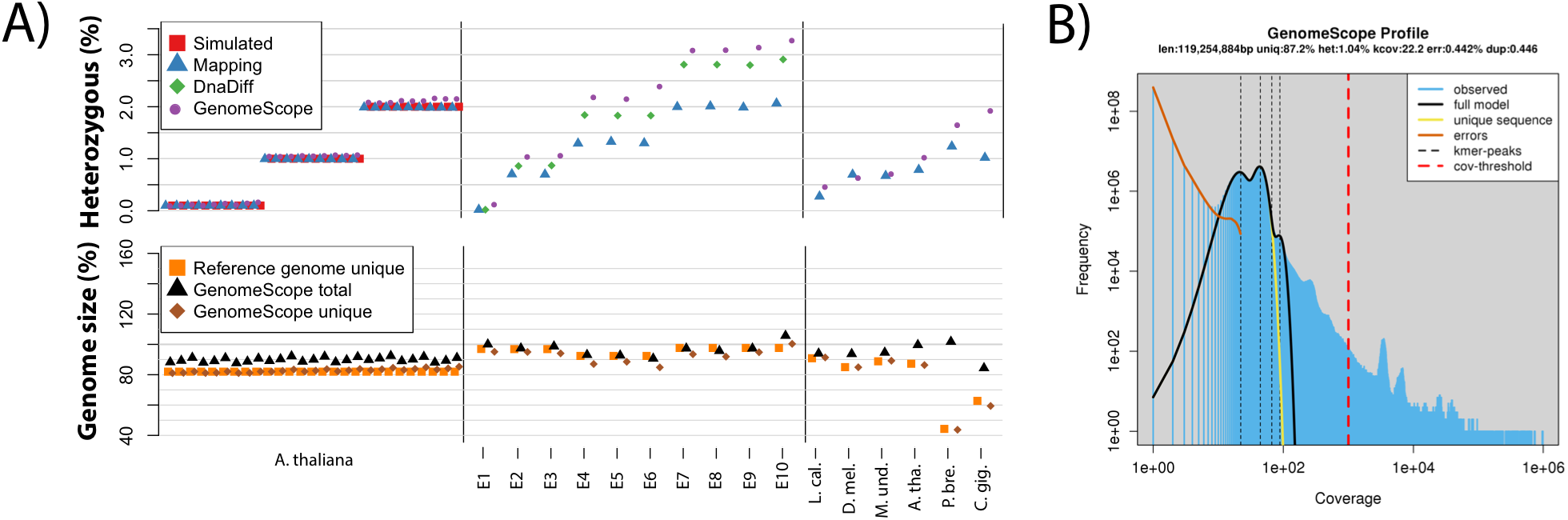
**(A)** *GenomeScope* heterozygosity, total genome size, and unique genome size estimates: (left) twenty seven simulated *A. thaliana* datasets with vary amounts of heterozygosity, sequencing error or read duplications; (middle) ten synthetic mixtures of real *E. coli* sequencing data; and (right) six genuine plant and animal sequencing datasets: *L. calcarifer* (Asian seabass), *D. melanogaster* (fruit fly), *M. undulates* (budgerigar), *A. thaliana* Col-Cvi F1 (thale cress), *P. bretschneideri* (pear), C. gigas (Pacific oyster). Also displayed are the true simulated values (Simulated), the results from a mapping and variant calling pipeline (Mapping), and a whole genome alignment (DnaDiff) where available. **(B)** GenomeScope k-mer profile plot of the *A. thaliana* dataset showing the fit of the GenomeScope model (black) to the observed k-mer frequencies (blue). The unusual peak of very high frequency k-mers (~10,000x coverage) were determined to be highly enriched for organelle sequences.

We next evaluated ten *E.coli* datasets where genuine sequencing reads from two divergent strains were synthetically mixed together (**Figure 1A**, middle). This allowed us to evaluate GenomeScope on real sequencing reads where the finished genome sequences, and hence their heterozygosity rates, could be precisely computed. We find high concordance to the results of the whole genome alignment of the reference genomes, although mapping the reads and calling variants resulted in artificially lower rates of heterozygosity because the short reads failed to map over the most heterozygous and repetitive regions. We also note that DnaDiff tends to underreport the rate of heterozygosity, especially if one genome is appreciably larger than the other, as it bases its estimate on those regions of the genomes that can be confidently aligned to each other while GenomeScope performs a more comprehensive genomewide analysis **(Supplementary Note 3)**.

Finally, we applied GenomeScope to six different genuine plant and animal data sets up to 1.1 Gbp in size with significant levels of heterozygosity and an assembled reference genome **(Supplementary Note 4)**. Since the available references were haploid, it was not possible to validate the results with whole genome alignment, but they were compared to the short read mapping results. The results are generally concordant, although the GenomeScope heterozygosity estimates were modestly higher than those from read mapping, similar to the *E. coli* results caused by short read mapping deficiencies, and most discrepant for the lowest quality draft genomes.

When examining real sequencing data, we introduced a parameter to exclude extremely high frequency *k-mers* (default: 1,000x or greater), since those often represented organelle sequences or spike-in sequences occurring hundreds to thousands of times per cell in *A. thaliana* that artificially inflated the genome size (**Figure 1b; Supplemental Note 1.3.2**). After accounting for the artificially high copy sequences, the inferred genome sizes of the real data sets were 99.7% accurate as confirmed by orthogonal technologies, such as the established reference genomes or flow cytometry when available **(Supplementary Note 4).**

## 4 Discussion

We have shown on 340 data sets that *GenomeScope* is a fast, reliable and accurate method to estimate the overall genome and read characteristics of data sets without a reference genome. Using the web application, users can upload their *k-mer* profile and seconds later *GenomeScope* will report the genomic properties and generate high quality figures and tables. As such, we expect *GenomeScope* to become a routine component of all future genome analysis projects.

For most genomes and for the experiments shown here, we recommend using *k=21,* as this length is sufficiently long that most *k-mers* are not repetitive but is short enough that the analysis will be more robust to sequencing errors. Extremely large (haploid size >>10GB) and/or very repetitive genomes may benefit from larger values of *k* to increase the number of unique *k-mers*. Accurate inferences requires a minimum amount of coverage, at least 25x coverage of the haploid genome or greater, otherwise the model fit will be poor or not converge **(Supplemental Note 2)**. GenomeScope also requires relatively low error rate sequencing, such as Illumina sequencing, so that most *k-mers* do not contain errors. For example, a 2% error rate is supported as it corresponds to an error only every 50bp on average, which is greater than the typical *k-mer* size used. However, raw single molecule sequencing reads from Oxford Nanopore or Pacific Biosciences, which currently average 5-20% error, are not supported as an error will occur on average every 5 to 20 bp and thus infer with nearly every *k-mer* (Goodwin, et al., 2016). Finally, GenomeScope is only appropriate for diploid genomes because the heterozygosity model it uses only considers the possibility for two alleles. In principle the analysis could be extended to higher levels of ploidy by considering additional peaks in the k-mer profile.

Future work remains to extend GenomeScope to support polyploid genomes and genomes that have non-uniform copy number of their chromosomes, such as aneuploid cancer genomes or even unequal numbers of sex chromosomes. In these scenarios the reported heterozygosity rate will represent the fraction of bases that are haploid (copy number 1) versus diploid (copy number 2) as well as any heterozygous positions in the other chromosomes. Addressing these conditions will require extending the *k-mer* model to higher copy number states.

## Acknowledgements

We would like to thank Arthur Delcher, Rachel Sherman, Steven Salzberg, and Junyan Song for their helpful discussions and testing. We would also like to thank the reviewers of the manuscript and all of the users of the system that provided helpful feedback and testing.

## Funding

NSF [DBI-1350041 and IOS-1237880]; NIH [R01-HG006677] to MCS.

*Conflict of Interest*: none declared.

## References

Bankevich, A., et al. SPAdes: a new genome assembly algorithm and its applications to single-cell sequencing. J Comput Biol 2012;19(5):455–477.

Bates, D.M. and Watts, D.G. Nonlinear Regression Analysis and Its Applications. John Wiley & Sons, Inc.; 1988.

Chikhi, R. and Medvedev, P. Informed and automated k-mer size selection for genome assembly. Bioinformatics 2014;30(1):31–37.

Gnerre, S., et al. High-quality draft assemblies of mammalian genomes from massively parallel sequence data. Proc Natl Acad Sci U S A 2011;108(4):1513–1518.

Goodwin, S., McPherson, J.D. and McCombie, W.R. Coming of age: ten years of next-generation sequencing technologies. Nat Rev Genet 2016;17(6):333–351.

Kajitani, R., et al. Efficient de novo assembly of highly heterozygous genomes from whole-genome shotgun short reads. Genome Res 2014;24(8):1384–1395.

Kelley, D.R., Schatz, M.C. and Salzberg, S.L. Quake: quality-aware detection and correction of sequencing errors. Genome Biol 2010;11(11):R116.

Li, H. Aligning sequence reads, clone sequences and assembly contigs with BWA-MEM. ArXiv e-prints 2013.

Li, H., et al. The Sequence Alignment/Map format and SAMtools. Bioinformatics 2009;25(16):2078–2079.

Li, X. and Waterman, M.S. Estimating the repeat structure and length of DNA sequences using L-tuples. Genome Res 2003;13(8):1916–1922.

Liu, B., et al. Estimation of genomic characteristics by analyzing k-mer frequency in de novo genome projects. 2013;arXiv:1308.2012.

Marcais, G. and Kingsford, C. A fast, lock-free approach for efficient parallel counting of occurrences of k-mers. Bioinformatics 2011;27(6):764–770.

Melsted, P. and Halldorsson, B.V. KmerStream: streaming algorithms for k-mer abundance estimation. Bioinformatics 2014;30(24):3541–3547.

Miller, C.A., et al. ReadDepth: a parallel R package for detecting copy number alterations from short sequencing reads. PLoS One 2011;6(1):e16327.

Phillippy, A.M., Schatz, M.C. and Pop, M. Genome assembly forensics: finding the elusive mis-assembly. Genome Biol 2008;9(3):R55.

Simpson, J.T. Exploring genome characteristics and sequence quality without a reference. Bioinformatics 2014;30(9):1228–1235.

Smolka, M., et al. Teaser: Individualized benchmarking and optimization of read mapping results for NGS data. Genome Biol 2015;16:235.

